# A Basal Ganglia Circuit Sufficient To Guide Birdsong Learning

**DOI:** 10.1101/222828

**Authors:** Lei Xiao, Gaurav Chattree, Francisco Garcia Oscos, Mou Cao, Matthew J. Wanat, Todd F. Roberts

## Abstract

Learning complex vocal behaviors, like speech and birdsong, is thought to rely on continued performance evaluation. Whether candidate performance evaluation circuits in the brain are sufficient to guide vocal learning is not known. Here, we test the sufficiency of VTA projections to the vocal basal ganglia (Area X) in singing zebra finches, a songbird species that learns to produce a complex and stereotyped multi-syllabic courtship song during development. We optogenetically manipulate VTA axon terminals in singing birds contingent on how the pitch of individual song syllables are naturally performed. We find that optical excitation and inhibition of VTA terminals have opponent effects on future performances of targeted song syllables and are each sufficient to reliably guide learned changes in song, consistent with positive and negative reinforcement of performance outcomes. These findings define a central role for reinforcement mechanisms in learning vocalizations and provide the first demonstration of minimal circuit elements for learning vocal behaviors.

## INTRODUCTION

The ability to imitate vocal behaviors is exceedingly rare, with humans, songbirds and parrots providing canonical examples of this remarkable trait^1–4^. Fluent production of learned vocalizations requires continual evaluation of performances using auditory-feedback. Neural signals indicating whether vocal performances are well and/or poorly performed relative to performance goals may provide general circuit mechanisms for maintaining and shaping vocalizations^5,6^. However, it is not known if evaluative signals indicating better performances or worse performances are sufficient to guide adaptive changes in vocal behaviors, or how such signals are implemented in the brain.

Zebra finches provide a useful model in which to test these questions because they learn a single courtship song during development and use extensive daily practice to maintain expert performance of this song in adulthood^3,7,8^. Courtship song is comprised of a series of short, spectrally distinct, syllables strung together in a stereotyped order. Typical song syllables are ~50 – 150ms long and the entire song motif contains 3 – 8 syllables. Learning and maintenance of song syllables depends on auditory feedback and is thought to involve trial-to-trial performance evaluation. Cortical^9–14^, basal ganglia^15–19^, and cerebellar circuits^20^ have all been uniquely implicated in song learning and neurons signaling aspects of performance evaluation have been identified in at least three regions in the songbird brain, including secondary and tertiary auditory cortical circuits^9,21^ and in the ventral tegmental area (VTA)^5^. Here we focus on the function of VTA neurons projecting to the striatopallidal vocal basal ganglia (Area X) because dopaminergic neurons in VTA have been broadly implicated in reinforcement learning^6,22,23^, and because lesion and pharmacological inactivation studies in songbirds indicate that the output of the vocal basal ganglia circuitry is important for initial song learning^14,17^ and continued vocal plasticity^15,24,25^. However, it is not known if signaling of better or worse performances alone is sufficient to guide changes in song, or if phasic changes in VTA-Area X activity can have opponent influences on song performances.

We applied axon-targeted optogenetic methods to excite or inhibit VTA axon terminals in the vocal basal ganglia of freely singing zebra finches in order to test the function of this circuit in vocal learning. On-line assessment of the fundamental frequency (pitch) of a targeted song-syllable permitted precise trial-to-trial targeting of optical manipulations based on natural variation in syllable pitch. We find that activation and inhibition are each sufficient to predictably guide changes in how song syllables are sung and that these manipulations guide rapid and reliable pitch learning in opposite directions. These findings define a minimal synaptic circuit for vocal learning and highlight unexpected precision in how the VTA-basal ganglia pathway can guide changes to vocal behaviors.

### Pitch Contingent Auditory Feedback Negatively Reinforces Learned Changes in Vocal Pitch

We first tested the ability of zebra finches to adaptively modify the pitch of a song syllable in a negative reinforcement learning task^10,19,26^. Zebra finches practice their song hundreds to thousands of times each day and exhibit a small amount of natural, trial-to-trial, variability in how they produce the pitch of individual syllables. Pitch-contingent auditory feedback (pCAF, Figures 1A–1F) targets white-noise playback when the pitch of a syllable is below or above an experimenter-defined threshold. Brief pulses of white-noise playback are thought to function as an aversive cue, perhaps perceived as an error in vocal performance^26^. We targeted 100ms pulses of white-noise playback (60–80 dB SPL) to low-pitch variants of individual song syllables in seven birds (single targeted syllable in each bird). Birds were acclimated to pCAF acoustic chambers for 3–4 days. On the first day of the experiment their song was continually recorded in order to establish baseline levels of the pitch for each syllable. Pitch-contingent auditory feedback was started in the morning of the second day. Birds rapidly learned to shift the pitch of the pCAF targeted syllable throughout the day, exhibiting significant increases in the pitch of the targeted syllable and significant decreases in the number of syllables that fell below the threshold for white noise playback (Figures 1E–1F). These findings confirm the ability of zebra finches to rapidly and selectively modify their song in a negative reinforcement task.

**Figure 1.**
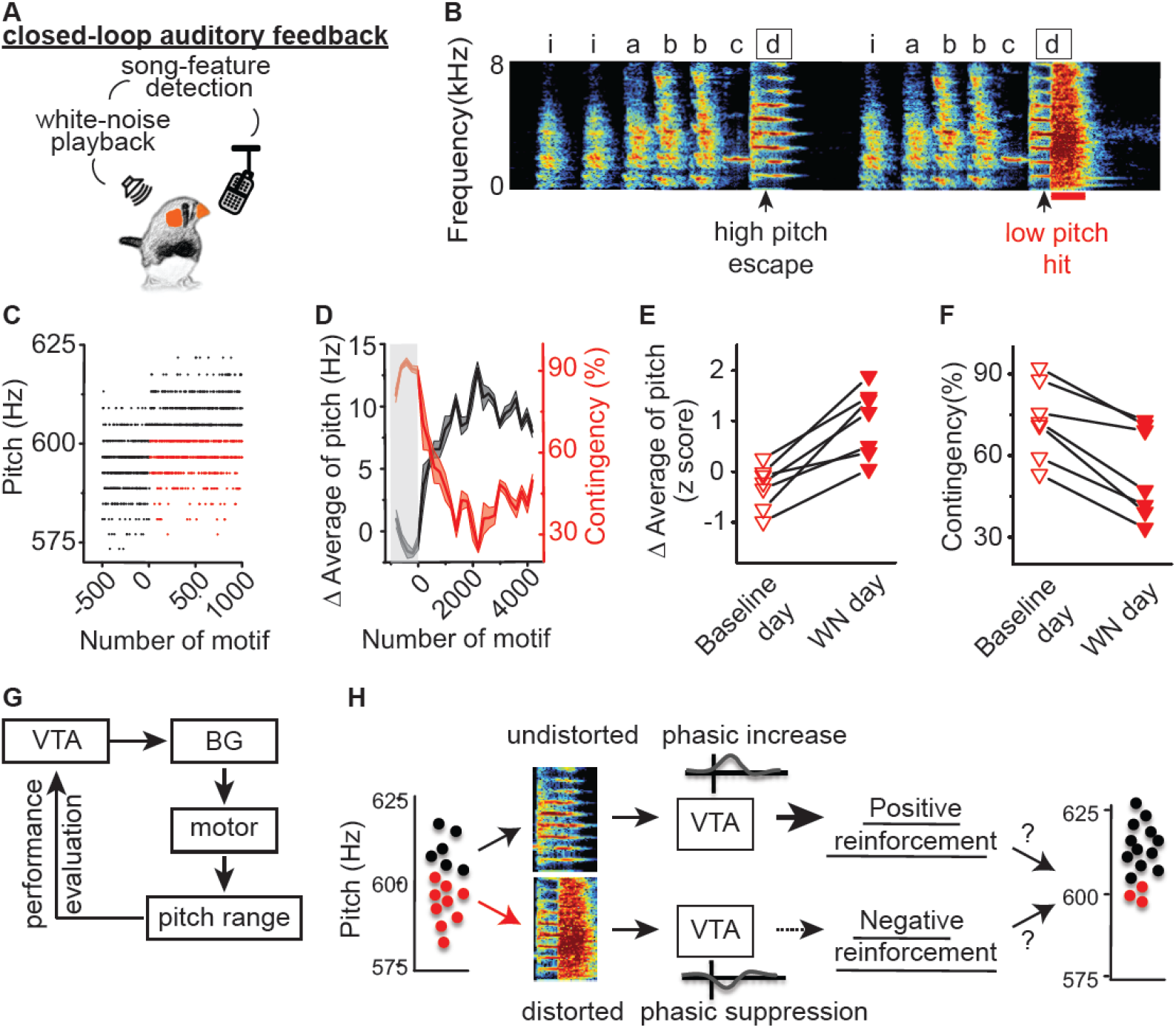
Pitch contingent auditory feedback guides pitch learning. **A)** Schematic of experimental design for close-looped pitch-contingent auditory feedback. **B)** Sonogram from the bird used in the pCAF experiment illustrated in figures 1C – 1D. White noise (WN) bursts were delivered over syllable ‘d’ during lower pitch variants. ‘abbcd’ indicate the syllables that comprise the bird’s motif. ‘i’ indicates an introductory note that usually occurs at the start of a song bout or before individual motifs. Arrowhead indicates a 5ms segment where the pitch of target syllable ‘d’ was measured; black box indicates target syllable. **C)** Plot of the pitch of syllable ‘d’ across 1,500 motifs before and during pCAF, each point corresponds to one rendition of the syllable. Closed-loop targeting of WN to lower pitch variants (red dots, ‘hit’) but not higher pitch variants (black dots, ‘escape’) resulted in an increase in the number ‘escape’ of trials. **D)** Plot of the running average of the pitch and hit rate (contingency) during the day of closed-loop pCAF illustrates the rapid increases in running average of pitch (black line) and concomitant decreases in contingency percentage (red line). Each point corresponds to a single syllable rendition and shaded region indicates ± one standard deviation; gray box indicates the baseline period before WN was delivered. **E)** Changes in running average of pitch during baseline day (open) and WN day (filled) in 7 birds in which WN was delivered to lower pitch variants (downward pointing triangles). WN delivering elicited increases in the running average of pitch (p=0.016, n=7, Wilcoxon matched-pairs signed-rank test). **F)** Changes in contingency percentage during baseline day (open) and WN day (filled) in 7 birds in which WN was delivered to lower pitch variants. WN elicited decreases in the contingency (p=0.015, n=7, Wilcoxon matched-pairs signed-rank test). **G)** Hypothetical source of instructive signal in natural pitch learning. **H)** General hypothesis tested in this paper: phasic increases and decreases in VTA neurons projecting to Area X encode positive and negative reinforcement signals that are each sufficient to guide song learning.

### Optogenetic Manipulation of VTA Terminals in the Songbird Vocal Basal Ganglia (Area X)

Learning to shift the pitch of an individual syllable embedded in a complex song motif is a form of motor skill learning that may depend on positive and/or negative reinforcement signals from VTA to Area X^5^ (Figures 1G–1H). Previous studies indicate that lesions to dopaminergic inputs to Area X largely block pitch learning during a negative reinforcement pCAF task^16^. To test if phasic increases or decreases in VTA_AX_ activity are sufficient to guide song learning, we sought to optogenetically manipulate VTA_AX_ axon terminals in freely singing birds. To enhance axonal distribution and membrane trafficking of virally expressed opsins, the intracellular targeting sequence of neurexin 1a was attached to the C-terminal end of channelrhodopsin (ChR2) or archaerhodopsin (ArchT3.0), resulting in viral constructs ChR2-nrxn-eYFP/2a-eYFP and ArchT-nrxn/2a-eYFP, referred to here as axChR2 and axArchT. Using retrograde tracing we found that VTA_AX_ neurons are predominantly located in the ventrolateral portions of VTA and that 88.1% of these neurons are dopamine (DA) positive, (Figure S1). Targeted viral injections into VTA revealed that we could efficiently infect dopaminergic VTA_AX_ neurons and robustly label their terminals in Area X (Figure 2A–2B, Figure S2).

**Figure 2.**
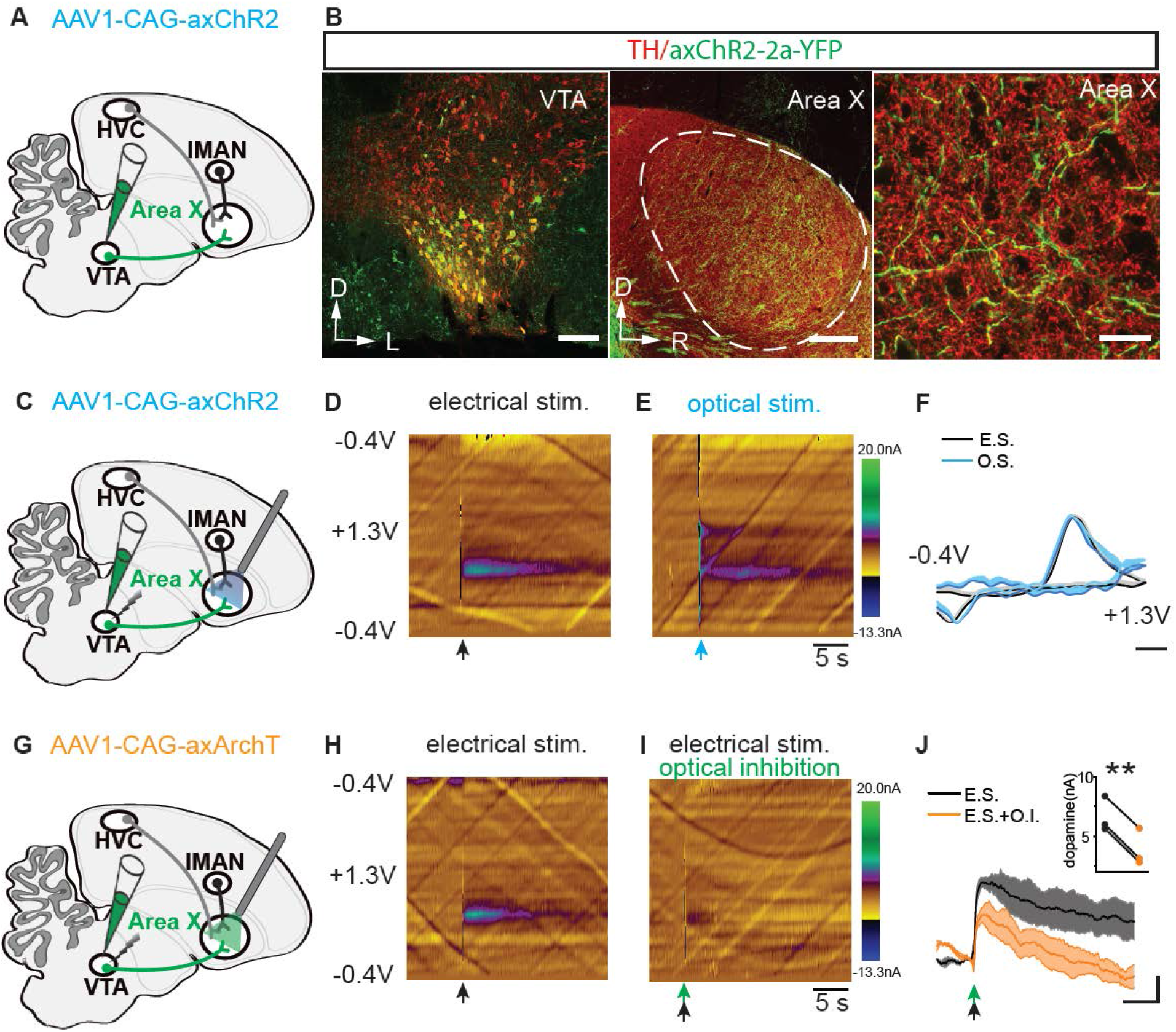
Optogenetic Manipulation of VTA Terminals in Singing Zebra Finches. **A)** Schematic showing injection of AAV1-CAG-axChR2 into. **B)** (Left image) Representative coronal section through VTA shows that most neurons infected with AAV1-CAG-axChR2-2a-YFP are TH positive and located in ventral and ventrolateral potions of VTA. Scale bar, 100 μm. (Middle image) Representative parasagittal section shows that Area X was well innervated with axonal terminals (green) arising from VTA. Dashed line circle outlines the border of Area X with darker staining of TH(red) relative to the surrounding basal ganglia region. Scale bar, 300 μm; D, Dorsal; L, lateral; R, rostral. (Right image) Enlarged image from Area X shows that axChR2 positive axons (green) are overlapping with TH staining (red). Scale bar, 20 μm. **C)** Schematic showing projection-specific optical stimulation of VTA terminals in Area X and electrical stimulation of VTA in axChR2+ birds. **D)** Representative voltammetric color plot of dopamine(DA) release in Area X following electrical stimulation of either VTA(n=2) or Area X(n=3) within parasagittal brain slices. Electrical stimulation of either VTA or Area X gave rise to comparable voltammetry profiles (data not shown). Black arrow, 50 – 100ms, 50 – 60hz of electrical stimulation. **E)** Representative voltammetric color plot of DA release in Area X following optical stimulation of VTA terminals in Area X within parasagittal brain slices of axChR2+ birds(n=5). Blue arrow, 100ms, 470nm. **F)** Background-subtracted cyclic voltammogram from electrical (E.S.) and optical (O.S.) stimulated DA release in the Area X, *ex vivo*. Single light pulse stimulation (O.S. blue, 100ms, n=5 birds) produced signature DA signals (E.S., black, 50 – 100ms, 50 – 60hz, n=5). Scale bar, 200 mV; shaded region indicates standard error of the mean. **G)** Schematic showing injection of AAV1-CAG-axArchT into VTA and projection-specific inhibition of VTA terminals in the Area X paired with electrical stimulation of VTA. **H)** Representative voltammetric color plots of dopamine(DA) release in Area X following electrical stimulation of VTA within a parasagittal brain slice of either control (n=2) or axArchT+(n=3) birds. Black arrow, 50ms, 60hz of electrical stimulation. **I)** Representative voltammetric color plots of DA release in Area X when electrical stimulation of VTA is paired with optical inhibition of VTA terminals in axArchT+ birds (n=3, green arrow, 100ms, 540nm). **J)** Averaged DA responses to electrical stimulation (E.S) of VTA with (black) or without(orange) optical inhibition (O.I.) of VTA terminals in the Area X of axArchT+ birds (n=3) as measured by FSCV. Scale bar, 2s, 2nA; shaded region indicates standard error of the mean. Insert panel, Optical inhibition resulted in reduced peak DA levels evoked by electrical stimulation (p=0.0002; paired t test).

We next tested our ability to optically manipulate axon terminals of VTA_AX_ neurons and the phasic release of DA. To test if optical activation or inhibition of axon terminals in Area X was sufficient control dopamine release, we made fast scanning cyclic voltammetric recordings from Area X several weeks following injection of either AAV1-axChR2 or AAV1-axArchT into VTA (Figure 2C–2J). We prepared brain slices maintaining axonal connections between VTA and Area X and used electrical stimulation of VTA somata to assess the electrochemical signature for dopamine in zebra finch Area X. We found that a 100ms pulse of blue (470nm) light was sufficient to reliably evoke dopamine release from VTA_AX_ axon terminals expressing axChR2 (recordings from 5 birds, Figure 2C-2F). In birds expressing axArchT we paired electrical stimulation of VTA somata with light inhibition of terminals in Area X in order to test the efficacy of axon-terminal optical inhibition. We found that a 100ms pulse of green (540nm) light was sufficient to significantly suppress electrically evoked dopamine release in Area X (Figure 2G–2J). Together, these findings indicate that brief optogenetic excitation or inhibition of VTA_AX_ axon terminals is sufficient to control phasic dopamine signaling in the songbird basal ganglia and open the door for testing the role of VTA_AX_ dopaminergic projections in shaping performance of learned vocal behaviors.

### Phasic Stimulation of VTA Axon Terminals Guides Bidirectional Learned Changes in Vocal Pitch

To test if manipulation of VTA_AX_ axon terminals is sufficient to guide learned changes in the pitch of a targeted syllable, we first used closed-loop, pitch-contingent optogenetic stimulation in freely singing adult zebra finches (100ms light pulse, 460nm, 3–5mW, n = 7 birds). Unlike pCAF experiments, these experiments do not rely on playback of an aversive auditory cue, but rather test if optogenetic manipulation alone is sufficient to guide learning (Figure 3A). We implanted adult male zebra finches with fiber optic cables overlying Area X 6–12 weeks after bilateral injections of axChR2 into VTA (Figures 3B–3C). Birds were allowed to acclimate for up to a week following implantation of fiber optic cables, as measured by a return in their normal daily singing behavior. We then targeted stimulation to natural syllable variants that fell within the lowest third of all pitch variants. On the first day of the experiment birds were continuously recorded to establish baseline pitch levels for the syllable to be targeted. The next morning, birds received optogenetic stimulation contingent on how they sang their syllable during individual song motifs. Illumination onset occurred within 25ms (24 ±0.4ms) of the syllable-pitch measurement and persisted for 100ms, a temporal window overlapping with the production of the targeted song syllable. Birds readily sang through optical pulses and did not appear to have any overt changes in their singing behavior (n = 7 birds, Figures 3D). Visual inspection of song recordings or videos of singing birds during optical stimulation did not reveal light-evoked truncations in song, altering of syllable ordering, syllable dropping, or other overt changes in song timing.

**Figure 3.**
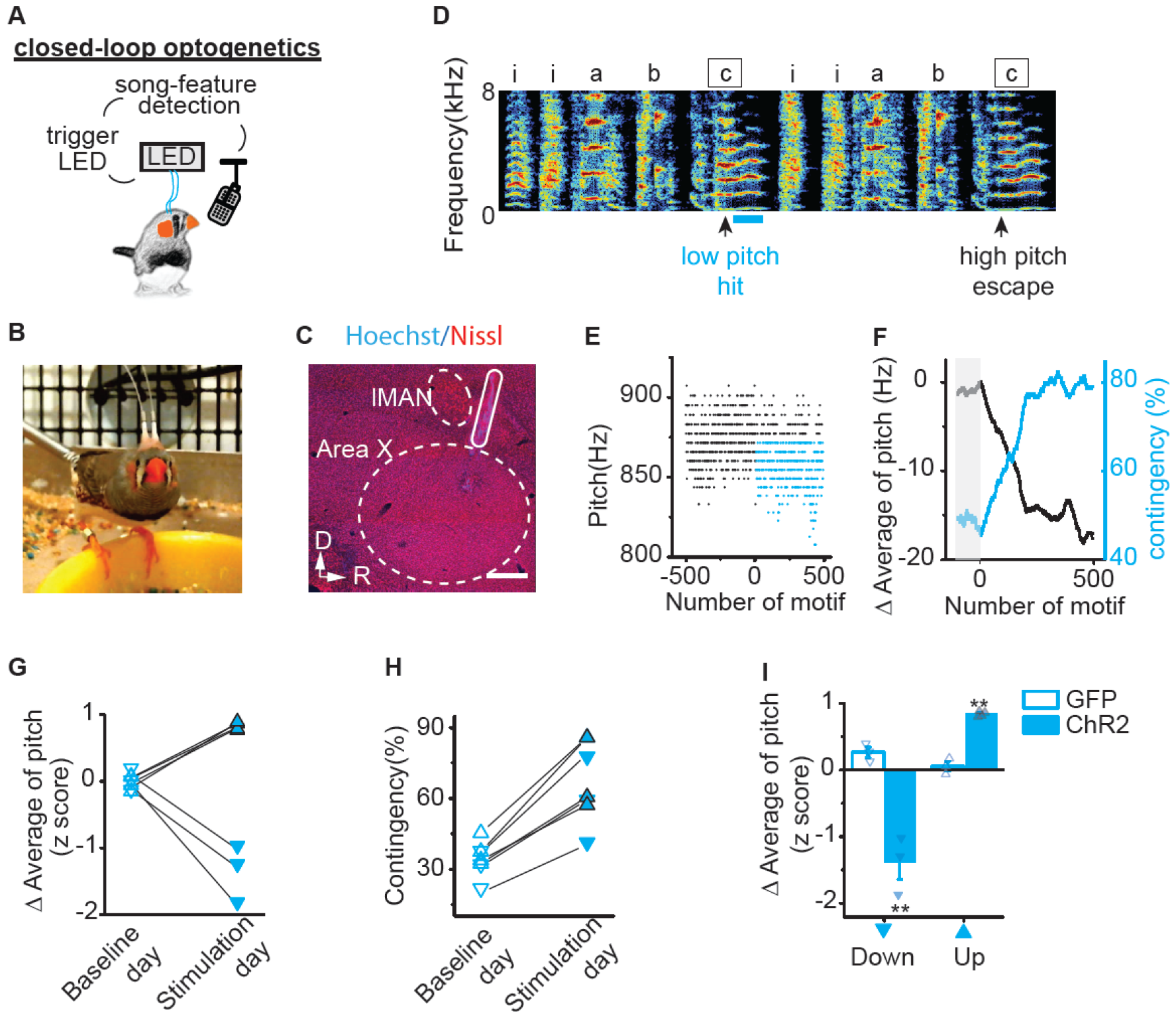
Pitch-Contingent Optical Excitation of VTA-Area X Terminals is Sufficient to Bidirectionally Guide Pitch Learning. **A)** Schematic of closed-loop pitch-contingent optogenetic experimental paradigm. **B)** A zebra finch with optical cannula bilaterally implanted over Area X. **C)** Parasagittal section shows implantation track of fiber optic overlying Area X and anterior to IMAN. Dashed lines outline both the border of Area X and IMAN. Scale bar, 300 μm. **D)** Sonogram from the bird used in the closed-loop optical stimulation experiment illustrated in figures 3E – 3F. Light pulses (~460 nm,100ms) were delivered over syllable ‘c’ during lower pitch variants (hit) and not during higher pitch variants (escape). **E)** Plot of the pitch of syllable ‘c’ across 1,000 motifs before and during optical stimulation, each point corresponds to one rendition of the syllable. Closed-loop optical stimulation of target syllables ‘c’ to lower pitch variants (blue dots) but not higher pitch variants (black dots) resulted in an increase in the number of ‘hit’ trials. **F)** Plot of the running average of the pitch and hit rate (contingency) during the day of close-looped optical stimulation illustrates the rapid decreases in running average of pitch (black line) and concomitant increases in contingency percentage (blue line). Each point corresponds to one rendition of the syllable; gray box indicates the baseline period before optical stimulation. **G)** Changes in running average of pitch during baseline day (open) and stimulation day (filled) for experiments in which optical stimulation was delivered to variants with higher pitch (upward pointing triangles with black outline, n=4) or lower pitch (downward pointing triangles, n=3), resulting in significant upward or downward shift in pitch (p=0.016, n=7, Wilcoxon matched-pairs signed-rank test). Changes in running average of pitch are expressed in units of the standard deviation of the last baseline session (z score). **H)** Closed-loop optical stimulation of syllables with either higher pitch (upward pointing triangles with black outline, n=4) or lower pitch (downward pointing triangles, n=3) elicited increases in contingency (33.81 ±2.92% to 66.94 ±6.47%, p= 0.016, n=7, Wilcoxon matched-pairs signed-rank test). Contingency on baseline day (open) and stimulation day (filled) was determined according to the same preset threshold for each individual experiment. **I)** Closed-loop optical illumination of either higher pitch variants (up) or lower pitch variants(down) elicited upward (p=0.0045) or downward (p=0.015, Unpaired t test with Welch’s correction) shift in running average of pitch in axChR2 birds (filled, up n=4, down n=3) but not in GFP birds (open, up n=3, down n=3). Error bars indicate standard error of the mean.

We found rapid and reliable changes in the pitch of the targeted syllable following pitch-contingent optogenetic stimulation. For the experiments shown in figures 3D–3F, we targeted optogenetic stimulation only to those renditions in which the bird sang syllable ‘c’ with a pitch lower than 875 Hz (Figure 3D–3F). The pitch of syllable ‘c’ decreased by 17.1 Hz, 1.2 times the standard deviation of its baseline values, during the first day of stimulation (Figure 3F). Correspondingly, as the bird learned to shift the pitch of the targeted syllable, the number of syllables that reached threshold for light stimulation increased by a third, from 46% to 79%. Across experiments, we found that targeting optogenetic stimulation to syllable renditions in the lower third of the normal pitch range resulted in decreases in the average pitch of that syllable, consistent with stimulation functioning as a positive reinforcement signal (downward pointing triangles always correspond to illumination targeting lower pitch variants and upward pointing triangles correspond to illumination targeting higher pitch variants, Figure 3G). The running average of pitch decreased by 1.39 ±0.25 standard deviations during the stimulation day while it changed only 0.02 ±0.06 standard deviations during the baseline day (Figure 3G). The rapid decreases in syllable pitch observed following optogenetic stimulation to lower-pitch syllable renditions stand in sharp contrast to the rapid increases in pitch seen in our pCAF experiments (Figure 1C–1F). These results support the idea that phasic activation of VTA_AX_ axon terminals functions as a teaching signal that guides song learning by positively reinforcing associated performances.

Zebra finches are capable of increasing or decreasing the pitch of an individual song syllable in response to aversive reinforcement pCAF tasks^10,19^. To test if stimulation of VTA_AX_ axon terminals is sufficient to guide bidirectional changes in vocal performance we targeted light stimulation to syllable renditions sung within the highest third of all pitch variants, rather than in the lowest third. We found that targeting optogenetic stimulation to syllable renditions in the higher third of the normal pitch range resulted in substantial increases in the pitch of that syllable throughout the training day (upward pointing triangles, Figure 3G). The running average of pitch increased by 0.84 ±0.02 standard deviations during the stimulation day, while it changed only – 0.007 ±0.035 standard deviations during the baseline day (Figure 3G). These findings suggest that optogenetic stimulation of VTA_AX_ axon terminals can guide learned increases or decreases in syllable pitch and further support the idea that phasic activation of VTA_AX_ axon terminals is sufficient to positively reinforce associated performances. Consistent with this, as birds learned to shift the pitch of the targeted syllable during the stimulation day, the light stimulation contingency increased for all birds, regardless of targeting higher or lower pitch variants (Figure 3H).

Dopamine neurons are known to respond to salient or novel sensory events, and unexpected light flashes, if visible during singing, could possibly cue changes in behavior. To test if changes in syllable pitch reflect a non-specific effect on song behavior we conducted identical pitch-contingent optical manipulations in birds injected in VTA with viral constructs only expressing GFP. Targeting light flashes to GFP expressing VTA_AX_ axon terminals during performance of either the higher or lower pitch syllable variants did not drive pitch learning and resulted in significantly smaller changes in pitch than optogenetic activation (Figure 3I). These results show that phasic activation of VTA inputs to Area X is sufficient to reliably guide rapid pitch learning, and support the idea that phasic increases in activity signal better than expected performances outcomes^5^.

### Inhibition of VTA Axon Terminals is Sufficient to Negatively Reinforce Changes in Vocal Pitch

We next tested whether pitch-contingent optical inhibition of VTA_AX_ axon terminals was sufficient to guide learned changes in song. We implanted adult male zebra finches with fiber optic cables overlying Area X 6–12 weeks after bilateral injections of axArchT into VTA. On the experimental day, we targeted inhibition to all but the highest or lowest pitch variants of an individual syllable in the birds’ polysyllabic song (upper or lower 60 – 90% of variants were yoked to optical inhibition, 100ms light pulse, 520nm, 1.5 – 4 mW, see Methods for description of contingencies). In contrast to optical activation of VTA_AX_ axon terminals, we found that optical inhibition resulted in rapid shifts in syllable pitch which mirrored those seen in birds exposed to a negative reinforcement pCAF task (Figures 4A–4C, 1C–D). For the experiments shown in figures 4A–4C, we targeted optogenetic inhibition to renditions in which the bird sang syllable ‘e’ with a pitch lower than 595 Hz (orange dots in Figure 4B). The pitch of syllable ‘e’ increased by 7.5 Hz, 0.72 times the standard deviation of its baseline values, over the course of a single experimental day and the stimulation contingency rate decreased by 19.5% (Figure 4C). Across experiments, we found that targeting optogenetic inhibition to lower pitch renditions resulted in rapid and substantial increases in the pitch of the targeted syllable. The running average of pitch increased by 1.08 ±0.15 standard deviations during the experimental day while it changed only 0.11 ±0.18 standard deviations during the baseline day (Figure 4D). Moreover, we found that optogenetic inhibition of VTA_AX_ axon terminals was sufficient to guide bidirectional changes in the pitch of song syllables in a manner consistent with negative reinforcement signals. Targeting inhibition to low pitch variants resulted in birds increasing the pitch of their song syllable, while targeting inhibition to high pitch variants resulted in birds decreasing the pitch of their song syllable (targeting high pitch variants, upward facing triangles: running average of pitch decreased by 1.36 ±0.2 standard deviations during the experimental day while it only changed 0.33 ±0.08 standard deviations during the baseline day, p=0.016, n=7, Wilcoxon matched-pairs signed-rank test, Figure 4D). Consistent with this, as birds learned to shift the pitch of the targeted syllable during the stimulation day, the light stimulation contingency decrease for all birds, regardless of targeting higher or lower pitch variants (Figure 4E). Lastly, we found that birds expressing only GFP did not exhibit any shifts in the pitch following identical pitch-contingent illumination over Area X and resulted in significantly smaller changes in pitch than optogenetic inhibition (Figure 4F).

**Figure 4.**
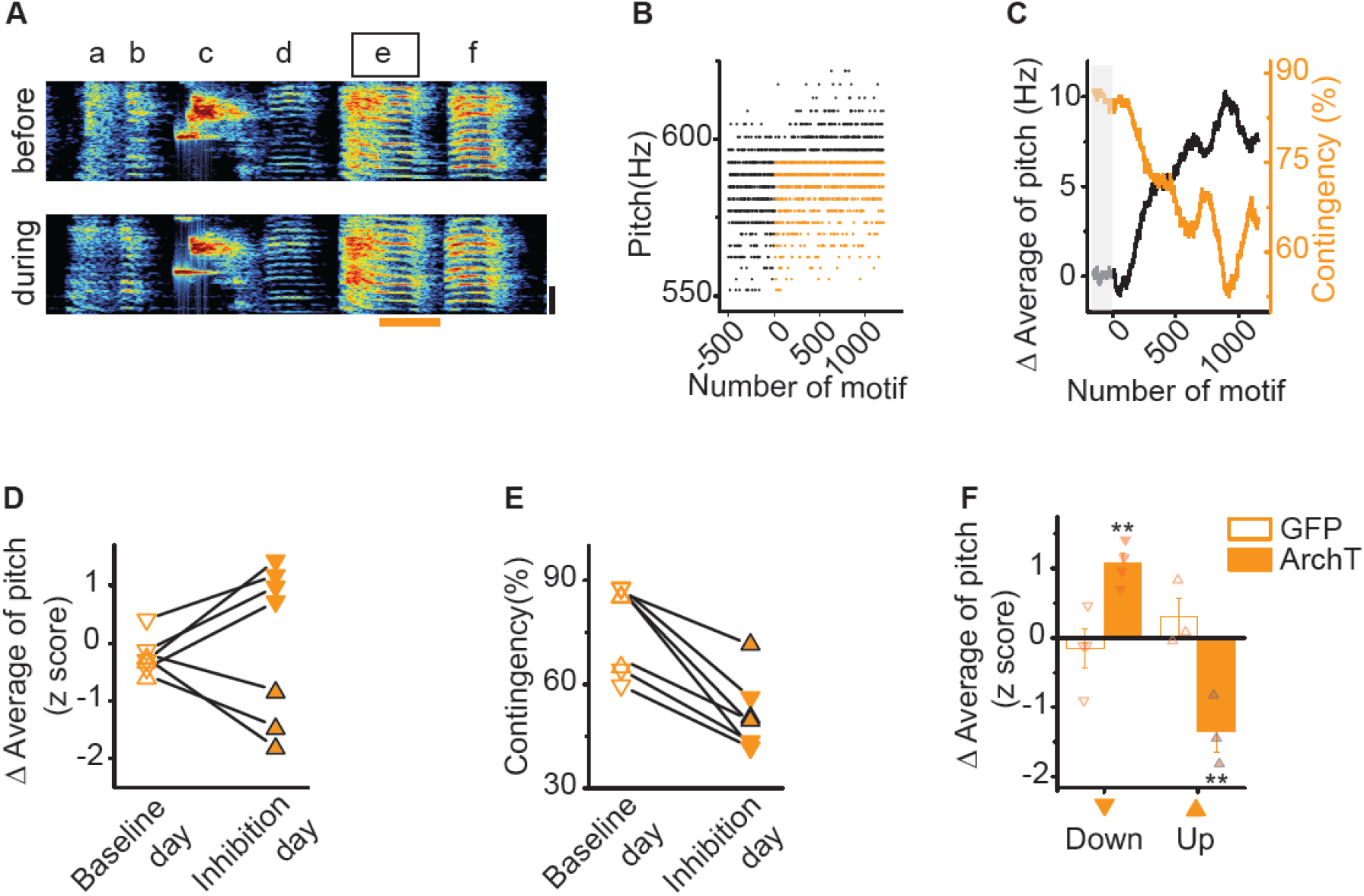
Pitch-Contingent Inhibition of VTA-Area X Terminals is Sufficient to Aversively Guide Pitch Learning. **A)** Sonogram from the bird used in the closed-loop optical inhibition experiment illustrated in figures 4B – 4C. Pitch contingent optical inhibition does not induce systematic changes in either song or syllable structure. Spectrograms of a song before (top) and during (bottom) an experiment in which light pulse (~520nm, 100ms, orange line) was delivered over syllable ‘e’ during lower pitch variants. **B)** Plot of the pitch of syllable ‘e’ across 1,701 motifs before and during optical inhibition, each point corresponds to one rendition of syllable. Closed-loop optical inhibition of target syllables ‘e’ to lower pitch variants (orange dots) but not higher pitch variants (black dots) resulted in an decrease in the number of ‘hit’ trials. **C)** Plot of the running average of the pitch and hit rate (contingency) during the day of close-looped optical inhibition illustrates the rapid increases in running average of pitch (black line) and concomitant decreases in contingency percentage (orange line). Each point corresponds to one rendition of the syllable; gray box indicates the baseline period before optical inhibition. **D)** Changes in running average of pitch during baseline day (open) and inhibition day (filled) for experiments in which optical inhibition was delivered to variants with higher pitch (upward pointing triangles with black outline, n=3) or lower pitch (downward pointing triangles, n=4), resulting in significant upward or downward shift in pitch (p=0.016, n=7, Wilcoxon matched-pairs signed-rank test). Changes in running average of pitch are expressed in units of the standard deviation of the last baseline session (z score). **E)** Closed-loop optical inhibition of syllables with either higher pitch (upward pointing triangles with black outline, n=3) or lower pitch (downward pointing triangles, n=4) elicited increases in contingency (p= 0.016, n=7, Wilcoxon matched-pairs signed-rank test). Contingency on baseline day (open) and stimulation day (filled) was determined according to the same preset threshold for each individual experiment. **F)** Closed-loop optical illumination of syllables with either higher pitch (up) or lower pitch (down) elicited downward (p=0.014) or upward (p=0.014, Unpaired t test with Welch’s correction) shift in running average of pitch in axArchT birds (filled, up n=3, down n=4) but not in GFP+ birds (open, up n=3, down n=4). Error bars indicate standard error of the mean.

These results demonstrate that optical inhibition of VTA_AX_ axon terminals is sufficient to guide changes in behavior consistent with negative reinforcement of vocal performances. Together with our optogenetic stimulation results, these findings provide strong causal support for VTA_AX_ dopamine neurons encoding positive and negative reinforcement signals that are each sufficient to guide rapid and selective learned changes to song.

### Optogenetic Manipulation of VTA Axon Terminals Guides Changes in Future Performances of Song

Reinforcement signals could shape changes in song by directly influencing motor performances^27–30^ or through or through evaluation of performance outcomes^5^. If VTA_AX_ neurons encode evaluation of performance outcome, it stands to reason that these signals should not have direct effects on motor performance, but rather have incremental effects on future performances. We examined these ideas in several ways. First, if activation or inhibition had direct effects on syllable pitch, we would expect immediate and stable effects on syllable behavior. Instead, we found accumulation of changes over the course of the day, consistent with iterative learning from an instructive signal (Figure 3F, Figure 4C). Second, direct motor effects on syllable pitch should also result in an increase in syllable variability. We found that the coefficient of variation of syllable pitch was not altered by our phasic activation or inhibition of VTA_AX_ axon terminals (Figure 5A–5B). This finding indicates that activation or inhibition of a subset of pitch variants resulted in learned changes across the entire distribution of pitch variants and supports the idea that learning resulted from incremental effects on future performances. Third, phasic activation or inhibition of VTA_AX_ axon terminals could result in direct motor effects on non-targeted vocal parameters (parameters other than pitch). To test this, we quantified several syllable performance parameters from interleaved light illuminated and escape trials during the first day of stimulation or inhibition, including amplitude, duration, Weiner entropy (measure of the width and uniformity of the power spectrum), goodness of pitch (measure of harmonic pitch periodicity), frequency modulation and amplitude modulation. We were unable to detect differences in any of the quantified features during optogenetic manipulation of VTA_AX_ axon terminals (n = 14 birds, Figure 5C–5D). Together, these findings demonstrate that closed-loop activation of VTA_AX_ axon terminals does not have immediate effects on motor performance, but rather guides learning by biasing the pitch of future performances.

**Figure 5.**
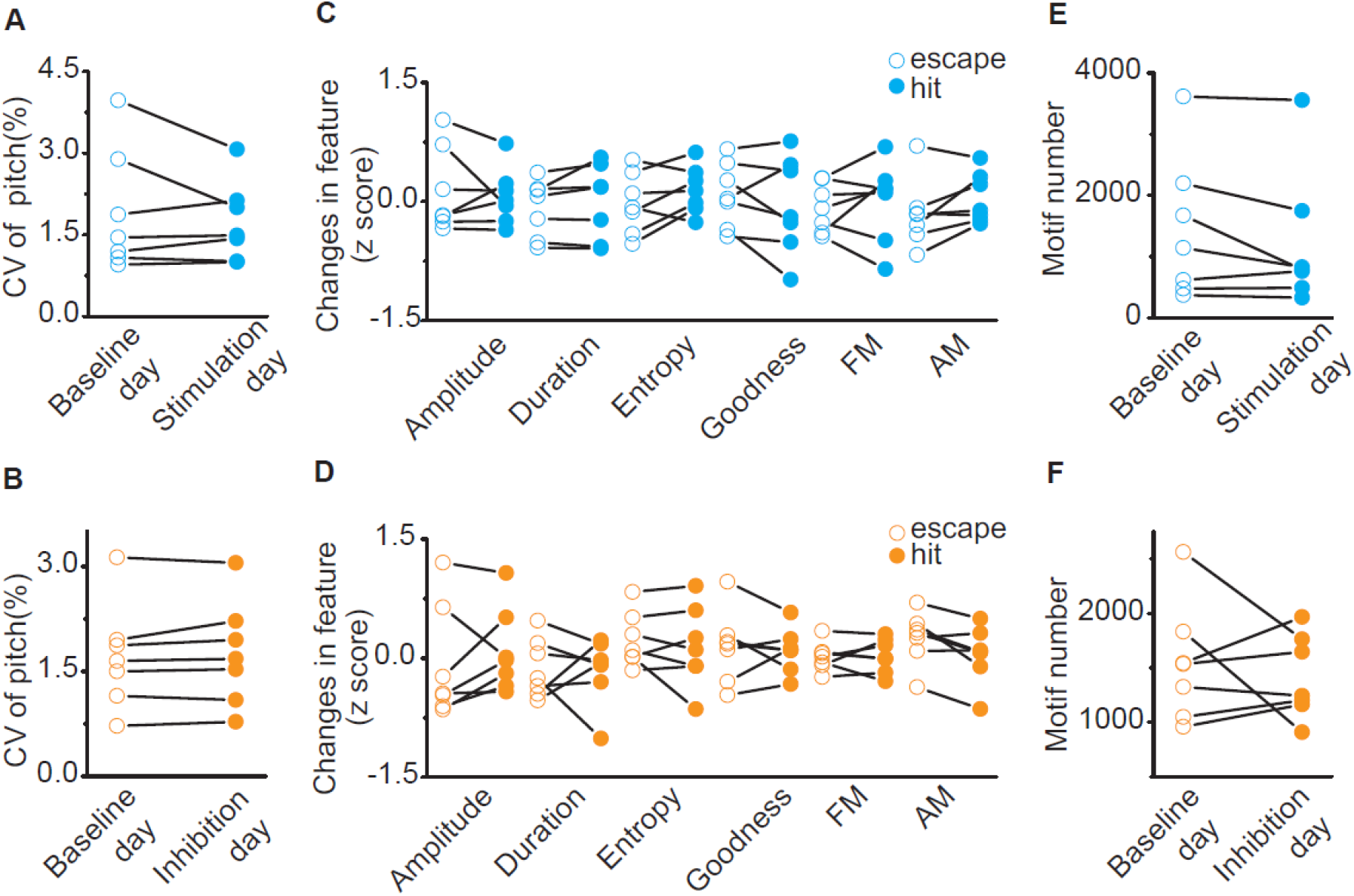
VTA-Area X Terminal Manipulations Do Not Have Motor or Motivational Effects on Song. **A)** Variability in pitch of target syllables for baseline day (open, CV = 1.91±0.42%) and stimulation day (filled, CV = 1.73±0.27%). Closed-loop optical stimulation of target syllables did not change the coefficient of variation of syllable pitch of target syllables (p=0.81, n=7, Wilcoxon matched-pairs signed-rank test). **B)** Variability in pitch of target syllables for baseline day (open, CV = 1.71±0.29%) and inhibition day (filled, CV = 1.75±0.28%). Closed-loop optical inhibition of target syllables did not change variability in pitch of target syllables (p=0.47, n=7, Wilcoxon matched-pairs signed-rank test). **C)** Spectral characteristics of ‘hit’ (filled) and ‘escape’ (open) syllable during the first stimulation session (200 motifs). Across experiments (n=7), there were no differences in amplitude(p=0.69), duration(p=0.30), Weiner entropy(p=0.16), goodness(p=0.38), frequency modulation (FM, p=0.69) and amplitude modulation (AM, p=0.11, Wilcoxon matched-pairs signed-rank test) between ‘hit’ and ‘escape’. **D)** Spectral characteristics of ‘hit’ (filled) and ‘escape’ (open) syllable during the first inhibition session (200 motifs). Across experiments (n=7), there were no differences in amplitude(p=0.38), duration(p=0.81), Weiner entropy(p=0.81), goodness(p=0.93), frequency modulation (FM, p>0.99) and amplitude modulation (AM, p=0.078, Wilcoxon matched-pairs signed-rank test) between ‘hit’ and ‘escape’. **E)** Motif number for baseline day (open, 1,444 ±441 motifs) and stimulation day (filled, 1,215 ±425 motifs). Closed-loop optical stimulation of target syllables didn’t change singing rate (p=0.15, n=7, Wilcoxon matched-pairs signed-rank test). **F)** Motif number for baseline day (open, 1,544 ±204 motifs) and inhibition day (filled, 1,413 ±144 motifs). Closed-loop optical inhibition of target syllables didn’t change singing rate (p>0. 999, n=7, Wilcoxon matched-pairs signed-rank test).

Beyond direct motor effects, repeated activation or inhibition of VTA_AX_ axon terminals could drive changes in pitch through changes in behavioral motivation. Dopamine signaling has been generally linked with reward, and successive activation or inhibition of VTA_AX_ axon terminals could lead to overall changes in the motivation to sing^31^. We examined singing rate in our birds and found that neither stimulation nor inhibition of VTA_AX_ axon terminals altered singing rate (Figure 5E–5F). We have shown that closed-loop optogenetic manipulation of VTA_AX_ axon terminals is sufficient to guide bidirectional changes in the future performances of song, independent of direct influences on ongoing song performance or generalized motivational changes in singing. These results indicate that phasic increases and decreases in dopaminergic input to Area X is sufficient to guide rapid and opponent changes in learned song, and support the idea that this single synaptic input provides a minimal circuit sufficient to direct vocal learning.

### VTA Axon Terminal Manipulations Drive Significant and Sustained Learned Changes in Song

The rapid, within-day changes in pitch, driven by optogenetic excitation or inhibition, reveal remarkable precision in the teaching signal that VTA conveys to Area X. Nonetheless, changes to syllable pitch in most of our birds remained within the natural range that the syllable could be produced prior to our optogenetic manipulations, raising concerns that manipulations of VTA_AX_ axon terminals are not sufficient to guide large scale changes in behavior, akin to those needed during initial learning of a new vocalization or recovery of vocal behaviors following peripheral or central injuries. To test if this teaching signal is capable of guiding sustained and large scale changes in vocal behavior, we extended our pitch-contingent optogenetic manipulations over several days, updating pitch-illumination thresholds each morning in order to continue driving changes in vocal behavior (3 – 12 days, n = 8 birds). Optogenetic stimulation and inhibition continued to have opponent effects on the direction of pitch learning. For example, successive targeting of lower pitch syllable renditions with optogenetic excitation resulted in cumulative decreases in syllable pitch, while optogenetic inhibition resulted in cumulative increases in syllable pitch over several days (Figure 6A). We found that changes in pitch learned on the first day of training were retained in the bird’s behavior the following morning, consistent with overnight consolidation of learned changes in song (**Figure S3**). Manipulations over several consecutive days, therefore, resulted in large changes in syllable pitch, with some birds shifting the pitch of their syllable by 3 – 6 standard deviations away from baseline values (quantified as z scores and absolute d-prime values (|*d*’|), see Methods, Figure 6A–E).

**Figure 6.**
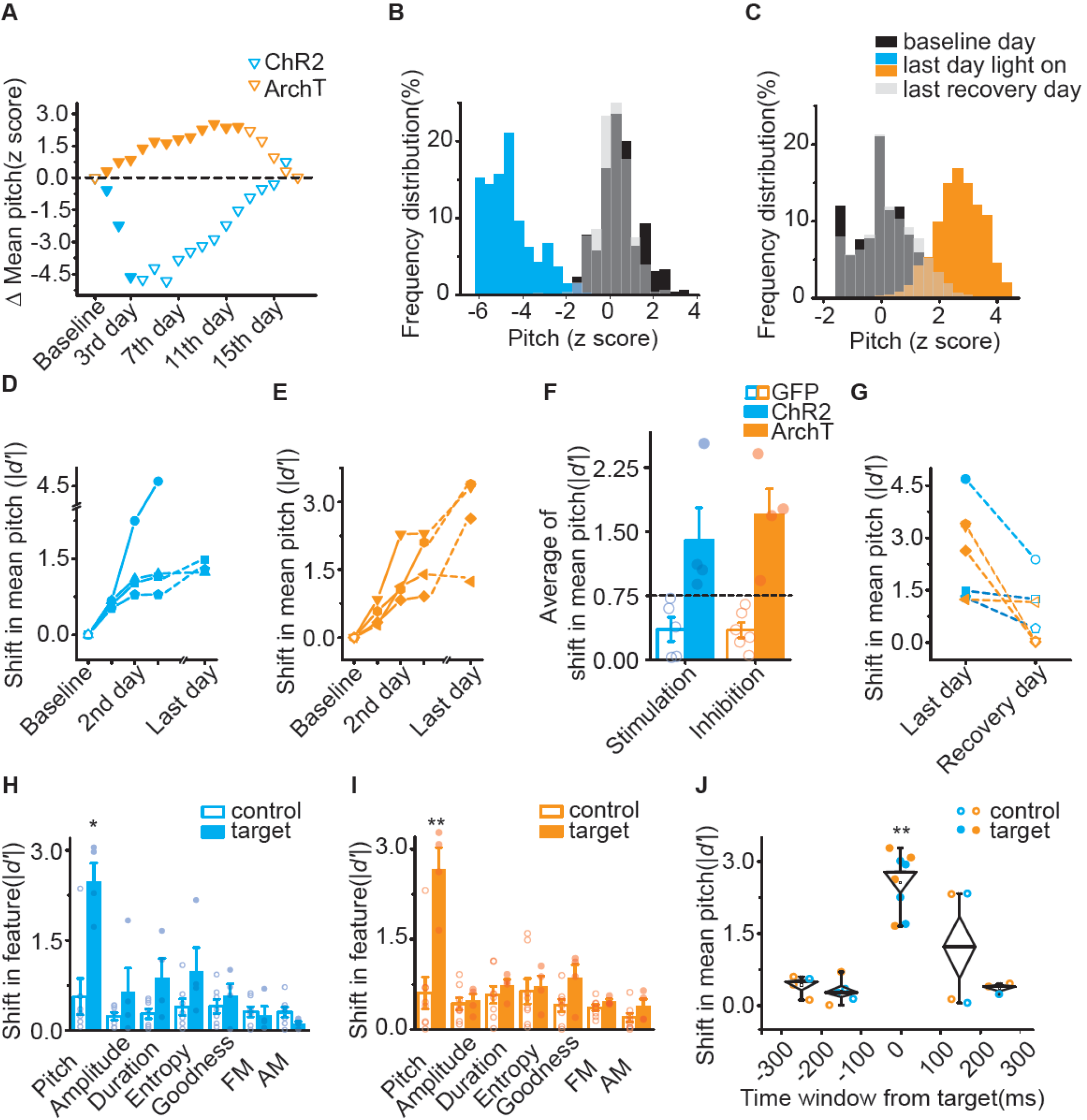
VTA-Area X Terminal Manipulations are Sufficient to Guide Opponent, Sustained and Selective Learned Changes in Song-Syllable Pitch. **A)** Difference in mean pitch between illumination day (filled) and baseline or recovery days (open) from one axChR2(blue) and one axArchT (orange) bird in which variants with lower pitch (downward pointing triangles) was targeted. Changes in mean pitch are expressed in units of the standard deviation of the baseline distribution(z score). **B)** Frequency distribution of pitch for the axChR2 bird shown in A. After illumination (blue), mean pitch shifted by 4.68 SD of baseline distribution(black) over 3 days and recovered (gray) to 0.37SD away from baseline distribution over 12 days. Pitch is expressed in units of the standard deviation of the baseline distribution (z-score). **C)** Frequency distribution of pitch for the axArchT bird shown in A. After illumination (orange), mean pitch shifted by 2.67 SD of baseline distribution(black) over 10 days and recovered (gray) to 0.02SD away from baseline distribution over 5 days. Pitch is expressed in units of the standard deviation of the baseline distribution (z-score). **D)** Shifts in mean pitch which expressed in unites of |*d*’| (see method) in 4 axChR2 birds over 3–6 days. E) Shifts in mean pitch which expressed in unites of |*d*’| in 4 axArchT birds over 5–10 days. **F)** Average of shift in mean pitch, expressed in units of |*d*’|, for GFP birds (n=6), axChR2 birds (n=4) and axArchT birds (n=4). Average shift in mean pitch for both axChR2 and axArchT birds were higher than 0.75, and also significantly higher than respective control GFP groups (axChR2, p=0.016; axArchT, p=0.0095; Mann-Whitney test). Error bars indicate standard error of the mean. **G)** Shifts in mean pitch for last illumination day and the day within a week following termination of illumination for both axChR2 (n=3) and axArchT (n=4) birds. Shifts in mean pitch were recovered toward to baseline within a week (changes in average of pitch Id’I, 2.6 ±0.5 vs 0.75 ±0.33, p=0.016, n=7, Wilcoxon matched-pairs signed-rank test). **H)** Shift in song spectral features for target (filled) and control (open) syllables of axChR2 birds(n=4). Changes in song were restricted to the pitch of target syllables (p=0.024, Mann-Whitney test). Spectral characteristics including amplitude (p=0.65), duration (p=0.11), Weiner entropy (p=0.11), goodness (p=0.53), FM (p=0.72), and AM (p=0.16) were not altered for either target or control syllables. Error bars indicate standard error of the mean. **I)** Shift in song spectral features for target (filled) and control (open) syllables of axArchT birds(n=4). Changes in song were restricted to pitch of target syllables (p=0.0081, Mann-Whitney test). Spectral characteristics including amplitude (p=0.68), duration (p=0.46), Weiner entropy (p=0.57), goodness (p=0.15), FM (p=0.21), and AM (p=0.15) were not altered for either target or control syllables. Error bars indicate standard error of the mean. **J)** Shift in mean pitch for target (filled) and control (open) syllables from both axChR2 birds (blue, n=4) and axArchT+ birds (orange, n=4) at millisecond time scale. Changes in pitch are restricted to target syllables (ANOVA, F_4,19_ = 13.62, P < 0.05, the diamonds denote s.e.m. and whiskers denote the 10–90% range).

To understand if these learned changes in pitch constitute behaviorally significant and sustained deviations, we assessed changes relative to a conservative threshold for naturally occurring variability of syllable pitch (|*d*’| = 0.75 significance threshold, Figure 6F)^32^. Both excitation and inhibition of VTA_AX_ axon terminals was sufficient to guide sustained and significant changes in syllable pitch, while similar manipulations in GFP birds did not result in learned changes in pitch (Figure 6F). In addition, we found that the standard deviation of syllable pitch at baseline positively correlated with maximum pitch shift birds were able to achieve (Figure S4). This finding held even for within-day changes in syllable pitch from birds receiving either activation or inhibition of VTA_vBG_ axon terminals. These results further underscore that our optogenetic manipulations do not alter syllable variability; rather changes in pitch are brought about by shifting the entire distribution of syllable variants over time.

Together, these results demonstrate that positive and negative reinforcement signals can guide large scale changes in the pitch of a song syllable, constrained only by the intrinsic variability in how the bird naturally sings the targeted syllable.

Although changes in performance, guided by optogenetic manipulation of VTA_AX_ axon terminals, appear to be consolidated in motor circuits, birds were still able to recover the baseline levels of syllable pitch once optogenetic manipulations were stopped. In 7 out of 8 cases in which we were able to monitor the recovery phase, the pitch of target syllables returned to its original range within 7 – 10 days after we ceased optogenetic manipulations, similar to recovery of normal song behavior at the end of pCAF training (Figure 6G)^10,19,26,32^. These results show that closed-loop manipulation of VTA_AX_ axon terminals is sufficient to guide large scale learned changes in song syllable behavior.

### Changes in Vocal Behavior are Spectrally and Temporally Precise

Fluent production of vocalizations involves the sequencing or concatenation of many small volitional movements. In birdsong this is reflected in the correct ordering of individual syllables and song notes, each with their own learned spectral and temporal features^33,34^. For reinforcement signaling from VTA_AX_ neurons to be a viable mechanism for learning and maintenance of song it should be able to guide changes in syllable performance that are both spectrally and temporally precise, and not result in changes to other features of a song syllable, such as its frequency modulation, or to changes in other portions of the song. To quantify this spectral and temporal precision, we focused on birds in which we optogenetically shifted the pitch of a song syllable for several days and confined our analysis to the day in which they displayed the largest shift in behavior, typically the last day of our optogenetic manipulations. All birds used in these experiments had at least two harmonic syllables in their core song motif, only one of which was targeted for pitch-contingent optogenetic stimulation or inhibition (n = 8 birds). We first asked if learned changes in pitch also resulted in changes to other features in the song syllable, or if they resulted in coincident changes to a non-targeted harmonic syllable in the song motif. We quantified several features of the song syllable and found that none of these other features were significantly changed when compared to baseline values, confirming that only the pitch of the targeted syllable had been modified in both our optogenetically stimulated and inhibited birds (Figure 6H – 6I). Additionally, we found that changes in the pitch of the target syllables did not result in any systematic changes in the features of the non-targeted harmonic syllables (control in Figure 6H – 6I). These findings show that positive and negative reinforcement signals are able to guide changes in a single spectral feature in a bird’s syllable, revealing remarkable precision in how these signals can influence learning of vocal behaviors.

Neuromodulatory signals, including dopaminergic signaling in the striatum, can act over relatively long timescales compared to the behavior of a single song syllable, which are typically only 50 – 150ms in duration. Indeed, dopamine signaling in the striatum is reported to mediate plasticity within a temporal window of 0.3 –2 seconds^35^. To examine the temporal precision of behavioral changes guided by optogenetic manipulation of VTA_AX_ terminals, we measured changes to the pitch of harmonic syllables produced immediately before or after the targeted syllable in the same birds described above. Measuring out from the time point in the target syllable when the fundamental frequency was calculated, we measured changes in the mean pitch of adjacent harmonic syllables produced between ±100 – 200ms and ±200 – 300ms before or after the target syllable. We found our effects on vocal pitch are largely restricted to the target syllables, which in these birds were ~100ms in duration (range = 47.19 – 148.9ms, mean = 96.16 ±37.2ms, Figure 6J) and tend to not extend to neighboring syllables as little as 100ms removed from the onset of illumination. In 6 of 8 birds we were unable to detect any changes to the pitch of harmonic syllables produced within ±100 – 200ms of the targeted syllable (7 of 9 syllables, Figure S5) and no changes to syllables produced within ±200 – 300ms of the targeted syllable (6 syllables). Together, these findings show that the VTA-Area X pathway can direct changes to an individual feature in a bird’s syllable and has the temporal resolution necessary to largely confine changes to only targeted syllables.

## DISCUSSION

Birdsong is perhaps the best studied naturally learned skilled motor behavior and the foremost model for investigating neural circuit mechanisms for learning vocalizations^1,3^. Like birdsong, many fine motor skills are learned through extensive practice and require continued training to maintain expert performance. Gaining proficiency in motor performance requires neural circuits capable of evaluating performance outcomes relative to motor goals and the ability to bias future performances in accordance with these evaluative signals. The present results define the VTA to basal ganglia pathway as a minimal circuit in the songbird brain capable of guiding song learning. We show that closed-loop activation or inhibition are sufficient to direct song learning and provide causal support for positive and negative reinforcement signals in learning vocalizations. Our findings demonstrate that VTA provides Area X with this instructive signal and that phasic manipulation of this signal during natural performances is sufficient to bidirectionally bias future performances of song. Models of basal ganglia-dependent reinforcement learning postulate that song learning relies on the convergence of three signals onto MSNs in the vocal basal ganglia^6^: a signal encoding information about the time-step in the song; a signal encoding information about motor variability (for example, if the pitch is going to be sung higher or lower at specific moment in the song); and a bidirectional reinforcement signal from VTA that functions as an eligibility trace for synaptic plasticity (see circuit schematics in Figure 2). Consistent with these models, studies have shown that cortical vocal basal ganglia pathways play an essential role in biasing future performances of song, that dopaminergic input is necessary for learning during negative reinforcement auditory feedback tasks, and that VTA_AX_ neurons exhibit phasic changes in activity during singing that are consistent with encoding of performance errors^5,16,19,24,25^. The present results link these findings with reinforcement learning models by providing the first causal evidence that phasic increases or decrease in VTA inputs have opponent effects on future song performances and are sufficient to guide song away from its natural performance range. The data presented here also extend beyond current models by highlighting an unappreciated temporal precision in reinforcement-learning, and the utility of reinforcement signals for learning fast sequential skilled behaviors. Our manipulations were sufficient to guide spectrally specific changes to song syllables as short as 50ms and confined changes in song to a 100–300ms time window. We suggest that this form of temporal precision is likely a common attribute of circuitry involved in evaluation of fine motor behaviors.

## METHODS

### Animals

All experiments were performed on adult male zebra finches (*Taeniopygia guttata*) that were raised in our breeding facility and housed with their parents until at least 50 days of age. During experiments, birds were housed individually in sound-attenuating recording chambers (Med associates) on a 12/12 h day/night schedule and were given ad libitum access to food and water. All procedures were performed in accordance with established protocols approved by the UT Southwestern Medical Center Animal Care and Use Committee.

### Viral Vectors

The recombinant AAV vectors were serotyped with AAV1 coat proteins and produced by the University of North Carolina vector core facility (Chapel Hill, NC, USA) with titer exceeding 10^12^ vg/ml. The self-complementary AAV(scAAV) vectors were serotyped with AAV1 or AAV9 coat proteins and produced by the Duke viral vector core facility (Durham, NC, USA) or in the lab with titer exceeding 5X10^11^ vg/ml. AAV1-CAG-ChR2(H134R)-nrxn-2a-EYFP or scAAV1-CBh-ChR2(H134R)-nrxn-EYFP viruses were used interchangeably for targeted stimulation of VTA_AX_ axon terminals, abbreviated to axChR2. AAV1-CAG-ArchT-nrxn-2a-EYFP or AAV1-CBh-ArchT-nrxn viruses were used interchangeably for targeted inhibition of VTA_AX_ axon terminals, abbreviated to axArchT. ScAAV1-Cbh-GFP virus was used as an opsin-negative control. Both scAAV1 and scAAV9-Cbh-EGFP were used for tracing experiments. All viral vectors were aliquoted and stored at −80 °C until use.

### Stereotaxic Surgery

#### Virus/tracer injection

All surgical procedures were performed under aseptic conditions. Birds were anesthetized using isoflurane inhalation (1.5–2%) and placed in a stereotaxic apparatus. Viral injections were performed using previously described procedures^10,11^ at the following approximate stereotaxic coordinates relative to interaural zero and the brain surface were (rostral, lateral, depth, in mm): Ov (2.8, 1.0, 5.75), the center of Ov was located and mapped based on its robust white noise responses; VTA relative to the center of Ov (+0.3, −0.2, +1.8)^9^; Area X (5.1, 1.6, 3.3) with 43-degree head angle or (5.8, 1.6, 3) with 20-degree head angle, the boundary of Area X was verified using extracellular electrophysiological recordings. For behavioral experiments, 0.7μl AAVs were injected into the VTA between 70–90dph after identification of target syllables and allowed at least 4–6 weeks for expression. For tracing experiments, 0.12μl Alexa Fluor 488–conjugated dextran amines (Invitrogen, CA, USA) or scAAVs were injected into the Area X or the VTA and allowed 3–5 days for sufficient retrograde transport and labelling.

#### Optical fiber placement

Birds were bilaterally implanted with optical fiber cannula (Prizmatix, Israel), prior to behavioral training and following the surgery procedure for viral injection. Fiber implants (200 or 250um, NA=0.66, Prizmatix) were targeted to the dorsomedial aspect of the Area X and were secured to the skull with C&B Metabond quick adhesive cement (Parkell Inc., NY, USA) followed by dental cement (Diamond Springs Inc., CA, USA). The optical fibers were connected to a LED source (λ = ~460 nm or ~520 nm; Prizmatix) via a rotary joint (Prizmatix) using an optic fiber sleeve (Prizmatix). LED power was adjusted to produce the desired output at the tip of the implanted optic probe (3–5mw for 460nm LED; 1.5–4mw for 520nm LED).

### Behavioral Assays

#### CAF program

Custom LabView software (National Instruments) was used for online detection of target syllables and implementation of optogenetic manipulation^19^. All birds used these experiments were pre-screened and only birds with well-defined harmonic syllables that permitted reliable detection included. Pitch was computed on a 5ms sound segment located 15–80ms into the target syllable. The target segment was constant for a given bird but varied between birds. Running average of pitch was calculated as the average of the pitch over the last 200 renditions (one session) of the target.

#### Habituation

Before baseline recordings, birds were given at least 1 week to recover from cannula implantation and habituate to singing with attached optical fibers. Songs were recorded for several days to measure baseline statistics on the pitch of targeted syllables and develop spectral templates to detect syllables in real-time and trigger closed-loop optogenetic manipulation. False positive and false negative rates were quantified and maintained under 10% for all birds.

#### Threshold setup and contingency calculation

Thresholds for triggering the LED light source were set at the running average of last baseline session for each animal, such that approximately one-third of baseline rage would be either supra-threshold or subthreshold. If the pitch met the escape criterion no illumination was triggered (‘escape’). Otherwise, a 100-ms light pulse was delivered within 25ms of the measurement (‘hit’). Contingency was calculated as the percentage of ‘hit’ renditions out of total rendition numbers for a day or a session. During baseline periods, contingency was calculated in the absence of illumination. For optogenetic activation experiments we targeted the upper or lower third of naturally produced pitch variants^27^. For optogenetic inhibition experiments we targeted the upper or lower two-thirds of pitch variants, similar to procedures used in white-noise playback experiments^10,19,24,26^.

#### Closed-loop optogenetic manipulation

The threshold for triggering illumination was constant during each day, and was updated each morning. Optogenetic manipulations were maintained over several days and not ceased until the pitch approached an asymptotic state within a day (Figure 6B, blue columns) or constant state for three consecutive days (Figure 6A, last three filled orange downward pointing triangles). The same training parameters were used for axChR2, axArchT and GFP experiments.

### Behavioral Analysis and Statistics

All behavioral events were recorded by computer systems. Data analysis for the pitch of targeted syllables were performed using custom software written in MATLAB (MathWorks; Natick,Massachusetts). To summarize effects across syllables, we expressed the daily changes in pitch of targeted syllables as z-scores:

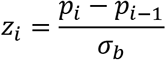

*p_i_* is the running average of pitch from last session on day *i* and *σ_b_* is the standard deviation of *last* baseline session.

*d*’ scores were computed to express the changes in mean daily pitch relative to last baseline day for each experimental day:

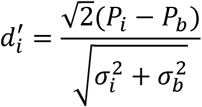

*P_i_* is the mean daily pitch on day *i* and 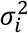 is variance. Day b refers to last baseline day.

In the case of equal variances 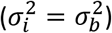, 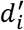 reports the changes in average between experimental day *i* and baseline day in the unit of SDs.

Maximum shift in mean daily pitch was computed as the difference in mean daily pitch between baseline day and the day d’ reaches maximum value (day_max_).

For analysis of spectral characteristics, Sound Analysis Pro (http://soundanalysispro.com/) was used to quantify longer segments (15–100 ms) of targeted or non-targeted harmonic stacks. Acoustic features were measured from one time matched session from baseline day and day_max_. d’ score for acoustics features were calculated using the same formula as for pitch shift.

For analysis of immediate motor effect, the first session of stimulation or inhibition day and time-matched session of last baseline day was analyzed. Z scores were computed for hit and escape renditions, using the last baseline session mean and standard deviation within each animal.

To determine whether parametric tests could be used, the Shapiro-Wilk Test was performed on all data as a test for normality. Unless otherwise noted, statistical significance was tested with non-parametric statistical tests; Wilcoxon signed-rank tests and Wilcoxon rank-sum tests were used where appropriate. Statistical significance refers to *P < 0.05, **P<0.02.

### Fast-Scan Cyclic Voltammetry

Birds were anesthetized using isoflurane and decapitated. The brain was quickly removed and immersed in ice-cold oxygenated zero-sodium ACSF containing the following (in mM): 225 sucrose, 3 KCl, 1.25 NaH2PO4, 26 NaHCO3, 10 D-(+)-glucose, 2 MgSO4, 2 CaCl2. The brain was then cut along the sagittal plane and the lateral side of the right or left hemisphere was glued onto a specimen tilting disc. The disc was tilted such that the vibratome blade entered the brain at a 10–15 ° angle. Slices (300 μm) containing Area X-Ventral Tegmental Area (VTA) were produced using a vibratome (Leica VT1200 / VT1200S) and an advancing speed of 0.12 mm/s. Slices were incubated in a custom-made holding chamber saturated with 95% / 5% O2/CO2 mix with reduced sodium ACSF containing the following (in mM): 60 NaCl, 75 sucrose, 2.5 KCl, 1.2 NaH2PO4, 30 NaHCO3, 25 D-(+)-glucose, 20 HEPES, 2 MgSO4, 2 CaCl2 at a temperature of 32°C for 40 min. The slices remained in the recovery chamber for at least another 40 min at room temperature before FSCV recording.

Slices were transferred to a recording chamber perfused with ACSF contained the following (in mM): 126 NaCl, 3 KCl, 1.25 NaH2PO4, 26 NaHCO3, 10 D-(+)-glucose, 2 MgSO4, 2 CaCl2 50 μM of L-DOPA, perfused at 3 ml min-1) at 31–33 C°. Stimulation of dopamine (DA) release was initiated typically 30 min after transfer to the chamber. Recordings were made in slices for up to 5 h after cutting.

Recordings were conducted using carbon-fiber electrodes (7 μm fiber diameter), the exposed carbon fiber tip was cut to a length of 30–150 μm. The tip of the CFE was gently lowered into the slice to a depth of 50–150 μm. The potential applied to the carbon fiber was ramped from −0.4 V (versus Ag/AgCl) to +1.3 V and back at a rate of 400 V/s during a voltammetric scan and held at −0.4 V between scans at a frequency of 10 Hz. All extracellular solutions were adjusted to 310 mOsm, pH 7.3–7.4, and aerated with a 95% / 5% O2/CO2 mix.

DA Release was evoked either by light emitted from a collimated light-emitting diode (470 nm) driven by a Cube LED Driver pE-300 (CoolLED) under the control of an Axon Digidata 1550B Data Acquisition System and Clampex 10.6 or through electrical stimulation. Light was delivered through the reflected light fluorescence illuminator port and the X 40 objective. For electrical stimulation a bipolar concentric stimulating electrode was placed in VTA controlled by a stimulus isolator (A365, WP) triggered by Axon Digidata 1550B data Acquisition System and Clampex 10.6.. Optimal stimulation employed a single 100ms light pulse. Electrical stimulation used a pulse-train at 50 Hz, 200–300 μA, 1 ms pulses, for 50 ms. After establishment of DA release, a laser light source (540nm; ~10mW) was used to illuminate Area X via the light fluorescence illuminator port and the X 40 objective for 100 ms single pulse.

### Histology

Immunohistochemistry experiments were performed following standard procedures. Briefly, birds were anesthetized with Ethanol (Virbac, TX, USA) and transcardially perfused with PBS, followed by 4% paraformaldehyde in PBS. Coronal sections (30μm) were cut using a freezing microtome (Leica SM 2010R, Leica). Sections were first washed in PBS, incubated in PBST (0.3% Triton X-100 in PBS) for 15min at room temperature (RT) and then washed with PBS. Next, sections were blocked in 5% donkey serum in PBST for 30 min at RT and then incubated with primary antibodies overnight at 4 °C. Sections were washed with PBS and incubated with fluorescent secondary antibodies at RT for 1 h. After washing with PBS, sections were mounted onto slides with Fluoromount-G (eBioscience, CA, USA). Images were acquired using a LSM 710 laser-scanning confocal microscope (Carl Zeiss, Germany) or an upright compound microscope (Leica DM5500 B, Leica). The primary antibodies used were: rabbit anti-tyrosine hydroxylase (AB152, Millipore, Germany), rabbit anti-GFP rabbit (A11122, Invitrogen, CA, USA) and mouse anti-GFP (A11120, Invitrogen, CA, USA). Primary antibodies were incubated with appropriate fluorophore-conjugated secondary antibodies (Life Technologies, Carlsbad, California, USA) depending on the desired fluorescence color.

### Data Availability

All data related to this research will be made available upon request.

### Code Availability

All analysis codes used in this research will be made available upon request.

### Author contributions

L.X., G.C. and T.F.R. conceived and designed all experiments. L.X. collected and analyzed the in vivo optogenetic behavioral, and the anatomical tracing data. G.C. collected and analyzed the pitch contingent auditory feedback data and optogenetic behavioral data. F.G.O. and T.F.R. collected the cyclic voltammetry data. M.C. analyzed and imaged the anatomical data and helped analyze optogenetic behavioral data. M.J.W. analyzed the voltammetry data and provided reagents and advice to help collect these data in songbirds. T.F.R. and L.X. wrote the manuscript. All authors read and commented on the manuscript.

## Acknowledgments

The authors thank Drs. Joseph Takahashi, Samuel Sober and Richard Hahnloser, as well as members of the Roberts laboratory, for discussions and/or comments on the manuscript, Dr. Jeremy Clark for advice and support in conducting fast scanning cyclic voltammetry experiments in songbirds, Dr. Claire Stelly and Merridee Lefner for fabrication of voltammetry electrodes, Drs. Karl Deisseroth and Edward Boyden for providing optogenetic constructs, Marguerita Kline for cloning of initial axon targeted constructs and Jennifer Holdway and Andrea Guerrero for laboratory support and animal husbandry. The Roberts laboratory was supported by grants from the National Science Foundation (IOS-1457206, IOS-1451034), the US National Institutes of Health (R01DC014364, R03MH111319), University of Texas BRAIN Initiative (362808, 362430), the Klingenstein-Simons Fellowship, and a NARSAD Young Investigator Grant (Essel Investigator) from the Brain & Behavior Research Foundation (T.F.R.).

## Author Information

The authors declare no competing financial interests. Correspondence and requests for materials should be addressed to T.F.R.

## Supplementary Figures and Legends

**Supplemental Figure 1.**
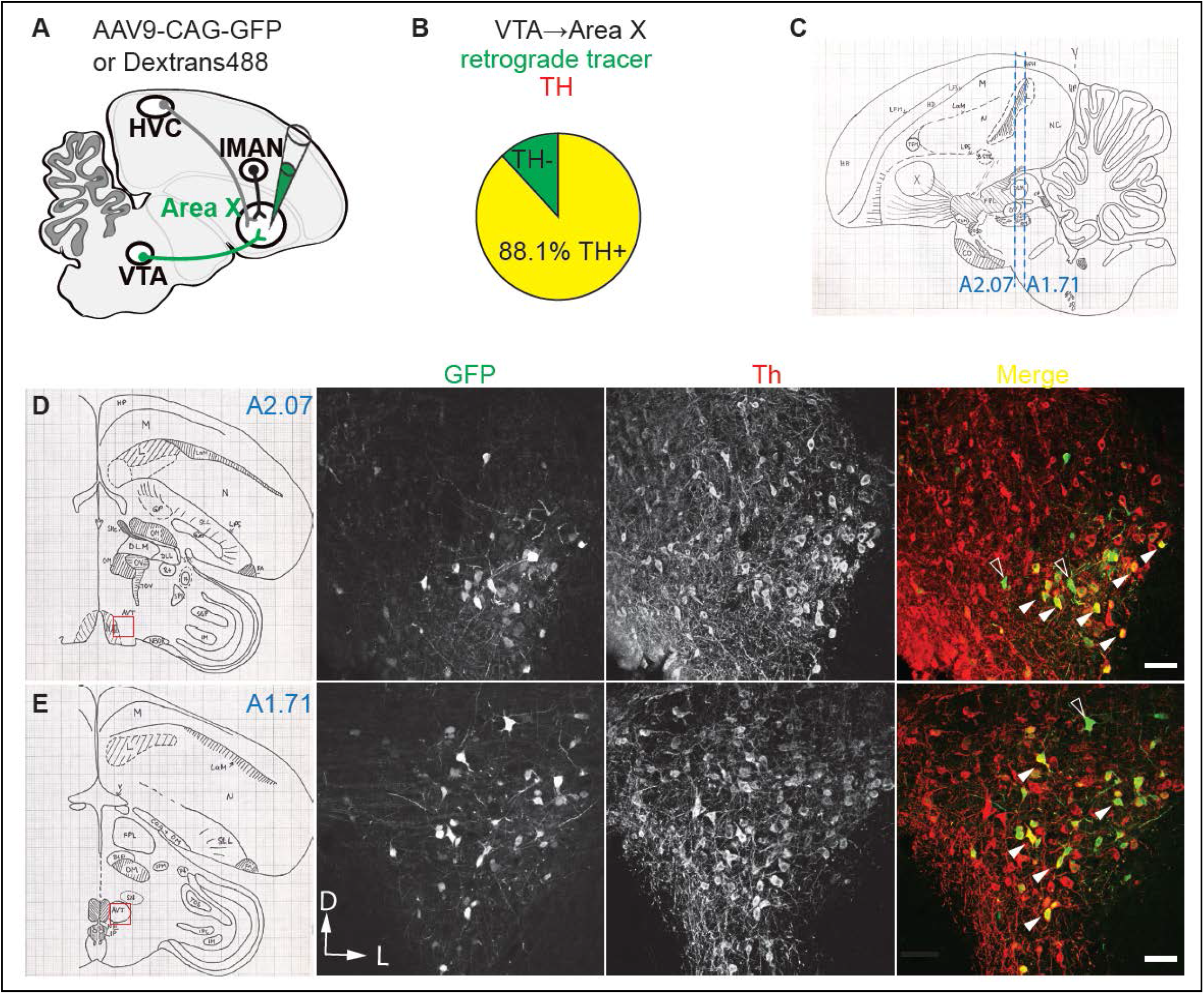
Anatomical identification of VTA_AX_ neurons. A. Schematic showing the design of retrograde-labeling experiments. Fluorescent retrograde tracers (dextran, Alexa Fluor, n=4) or AAV9-CAG-GFP(n=3) were injected into Area X. B. Retrograde-labeling experiments reveal that 88.1% of neurons (1926 out of 2185 in VTA) projecting into Area X area (yellow, filled arrowheads in figures D&E) are TH positive. C. Schematic representation of the cutting plane in sagittal section (cited from ‘a stereotaxic atlas of the brain of the zebra finch’). Blue dashed lines illustrate two transverse sections 2.07mm (A2.07, figure D) and 1.71mm (A1.71, Figure E) rostral of Y-point. D. and E. Confocal images taken from rostral VTA (figure D, A2.07) and caudal VTA (figure E, A1.71). Most of neurons projecting to Area X, located in the ventrolateral VTA, are TH positive (yellow, filled arrowheads), open arrowheads indicate TH negative Area X projecting neurons(green). Scale bar, 100 μm; D, Dorsal; L, lateral.

**Supplemental Figure 2.**
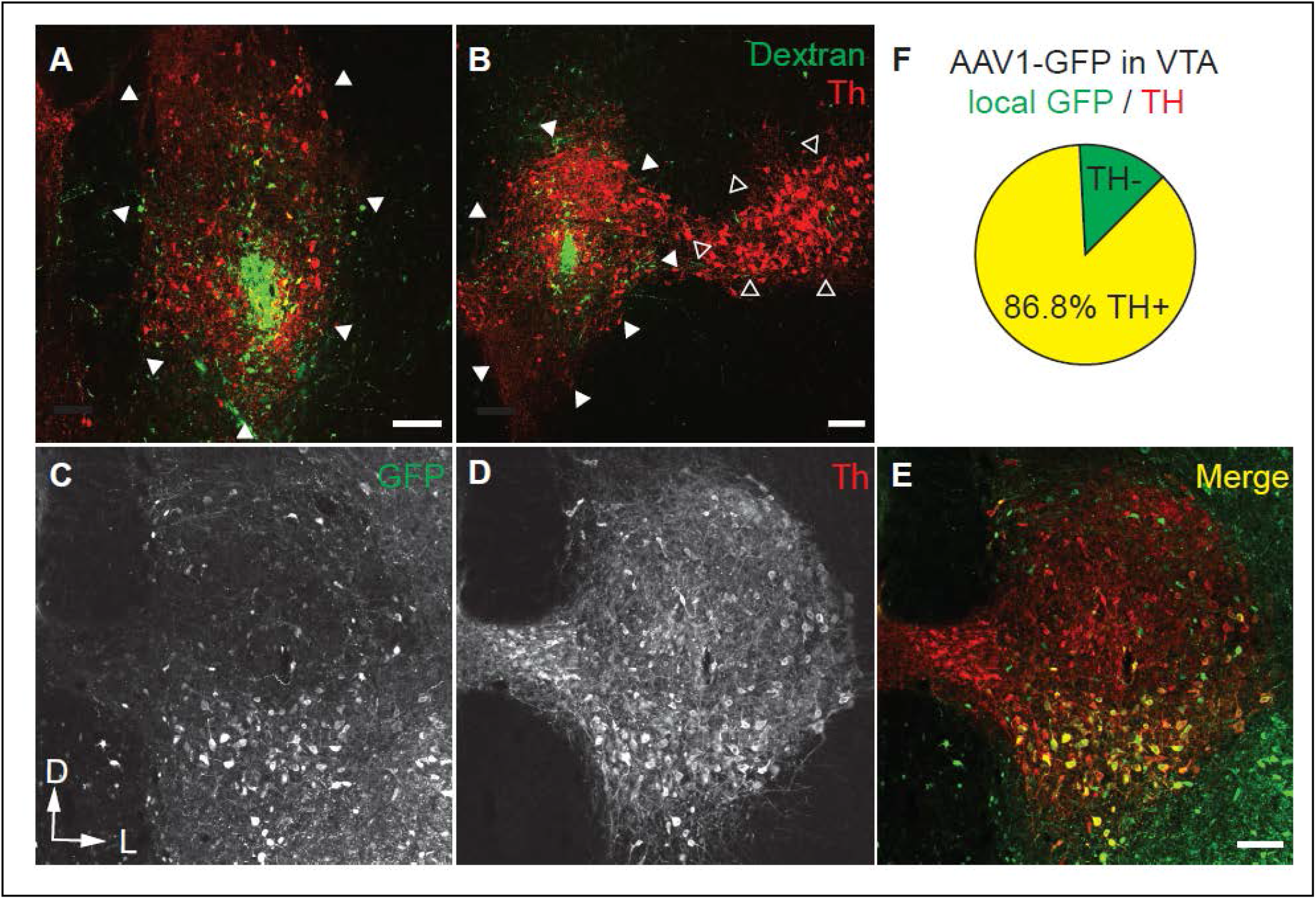
Targeting injections to VTA. A. and B. Representative coronal sections taken from a bird injected with fluorescent tracer (dextran 488) injected into VTA illustrates that our approach provides reliable targeting of VTA (figure A = A2.07 and figure B = A1.71). Filled triangles outline the border of VTA and open triangles outline the border of SNc. Scale bar, 100 μm. C. –E. Representative coronal sections taken from a bird with AAV1-CAG-GFP directly injected into VTA reveals that most of GFP positive neurons in VTA are TH positive. Scale bar, 100 μm. F. Injections of AAV1-CAG-GFP into VTA reveal that 86.8% of virally infected neurons (992 out of 1143, 8 hemispheres from 4 birds) in VTA are TH positive.

**Supplemental Figure 3.**
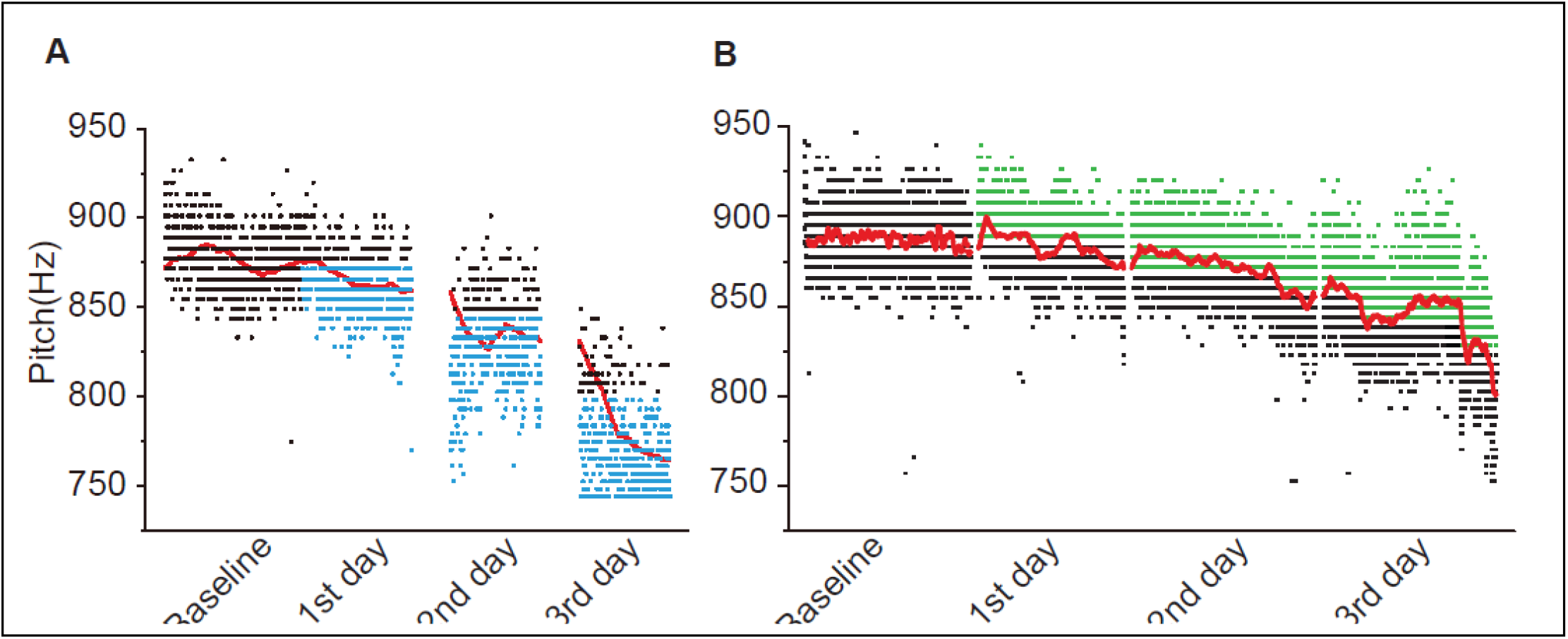
Raw pitch changes over 3 days for an axChR2 and a axArchT bird. **A-B** Representative plot of all pitch variants for one axChR2+bird with illumination to variants with lower pitch (Figure A, blue dot) and one axArchT+ bird with illumination to variants with high pitch (Figure B, green dot). During three consecutive days, both birds exhibited continued decrease in running average of pitch (red line). Black dot, baseline or ‘escape’ rendition.

**Supplemental Figure 4.**
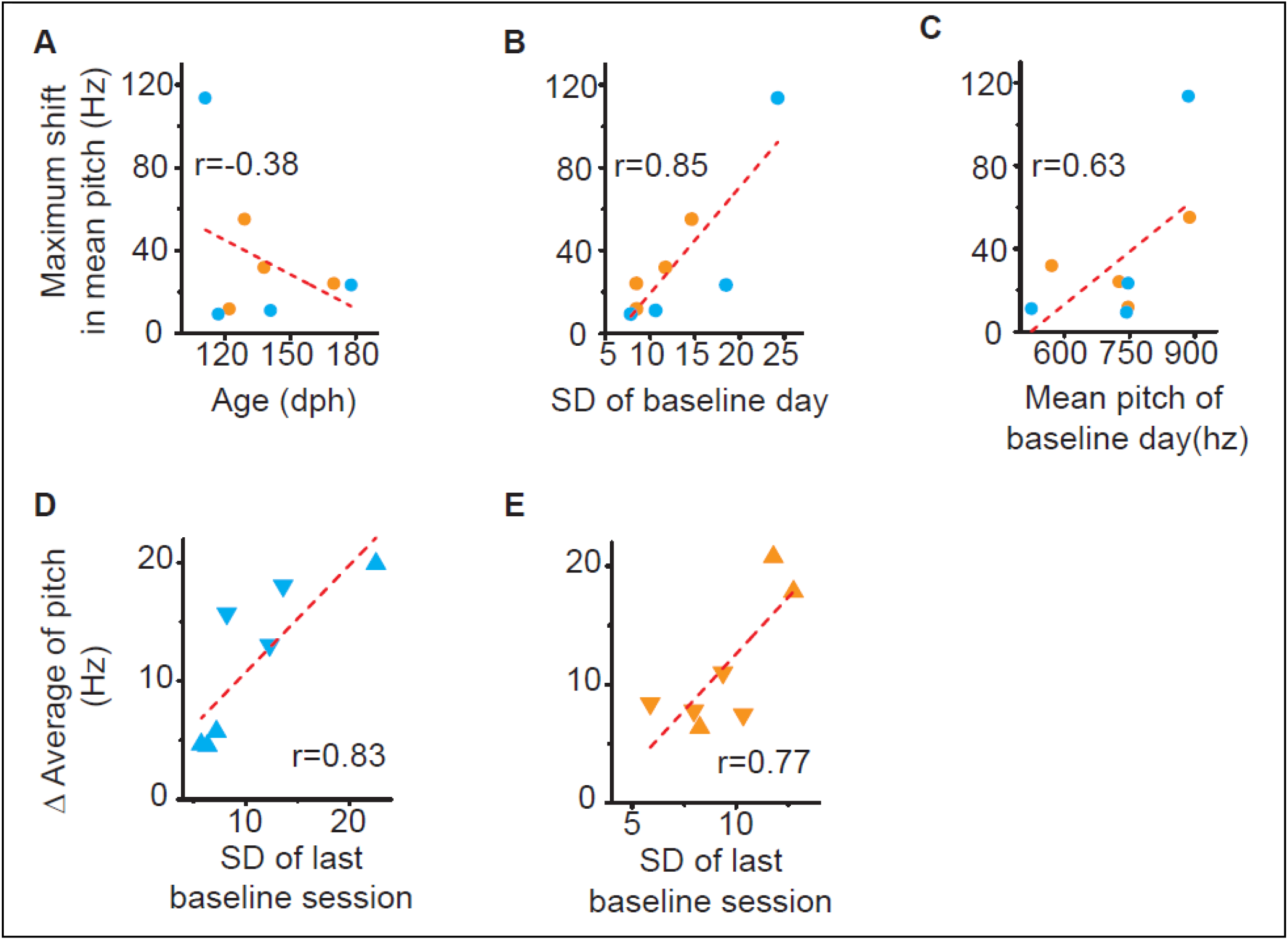
Optogenetic changes in pitch correlate with natural variation in syllable pitch. **A-C** Relationship between the magnitude of maximum shift in mean pitch and age of the bird or features of their target syllable. Across excitation and inhibition experiments, there was a positive correlation between the maximum shift in mean pitch and the SD of pitch on baseline day (**B**, r=0.85; p=0.0081, n=8, pearson). In contrast, the magnitude of maximal shift is not related with either age of the bird (**A**, r=−0.38, p=0.35, pearson) or mean pitch of target syllable (**C**, r=0.63, p=0.096, pearson). **D-E** Relationship between the standard deviation(SD) of pitch in last baseline session and the magnitude of shift in running average over the course of the stimulation day **(D)** or inhibition day **(E)**. Across experiments, there was a positive correlation between the SD of pitch in last baseline session and the magnitude of shift during both stimulation day (r=0.83, p = 0.020, Pearson) and inhibition day (r=0.77 p = 0.041, Pearson).

**Supplemental Figure 5.**
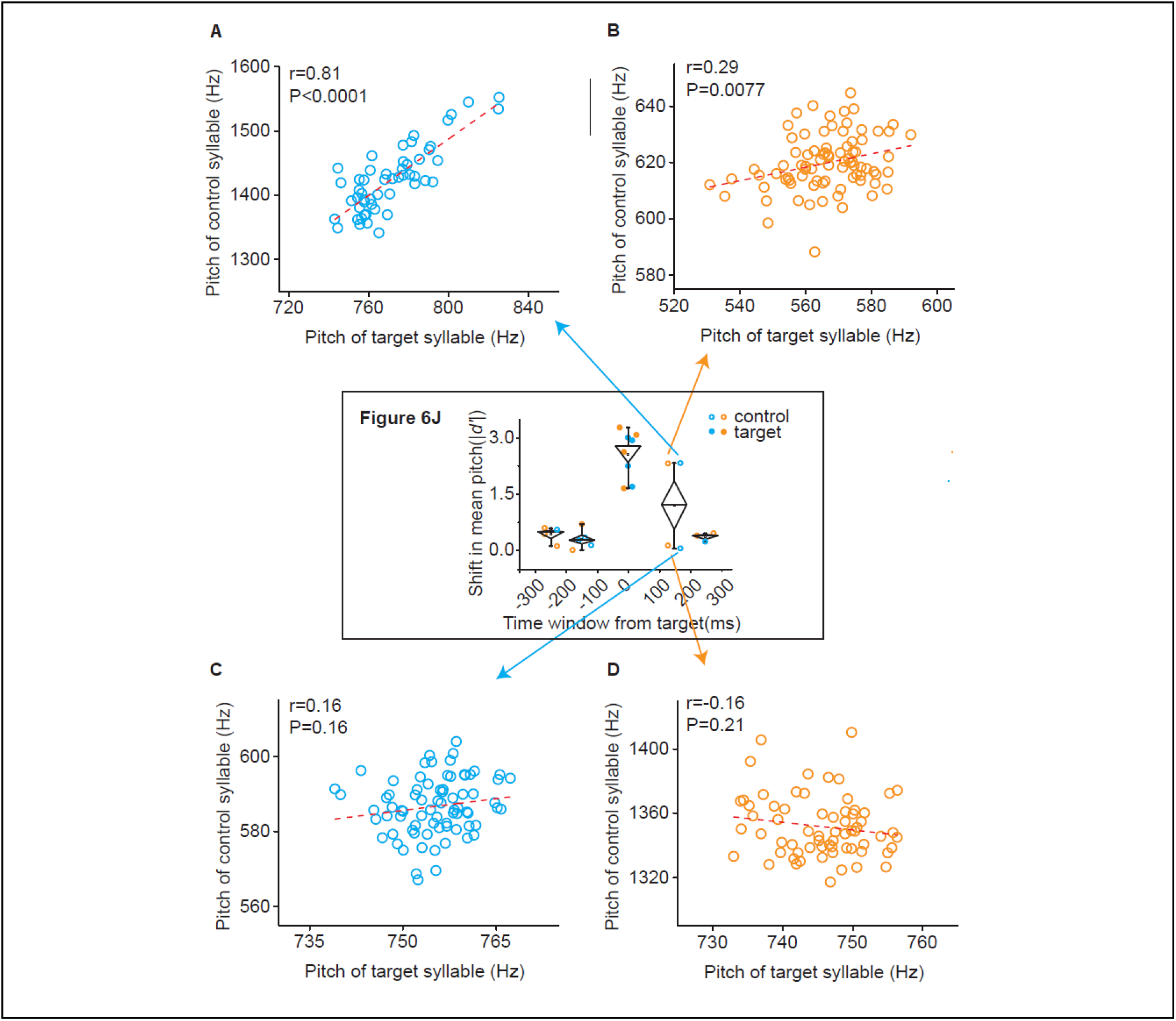
Baseline correlations in pitch between target syllables and control syllables within +100–200 window from 4 individual birds plotted in Figure 6J. This figure illustrates pitch correlations on baseline day for the four birds plotted in main Figure 6J. This analysis reveals that in the two birds in which the pitch of the control syllable shifted following optogenetic manipulations a positive correlation between the pitch of targeted and control syllables existed prior to optogenetic manipulations. For one axChR2+ bird and one axArchT+ bird in which pitch of control syllables shifted (Figure 6J, Blue, axChR2+, d’=2.33; orange, axArchT+, d’=2.32), the pitch of target syllable is positive correlated with the pitch of control syllable on baseline day (**S5 Figure A**, axChR2+, r=0.81, p<0.0001; **S5 Figure B**, axArchT+, r=0.29, p=0.0077). In contrast, the other axChR2+ bird and axArchT+ birds in which pitch of control syllables were not changed (Figure 6J, Blue, axChR2+, d’=0.056; orange, axArchT+, d’=0.13), there is no correlation between the pitch of target syllable and control syllable on baseline day (S5 Figure C, axChR2+, r=0.16, p=0.16; S5 Figure D, axArchT+, r=−0.16, p=0.21).

## References

1 Konopka, G. & Roberts, T. F. Insights into the Neural and Genetic Basis of Vocal Communication. Cell 164, 1269–1276, doi:10.1016/j.cell.2016.02.039 (2016).

2 Petkov, C. I. & Jarvis, E. D. Birds, primates, and spoken language origins: behavioral phenotypes and neurobiological substrates. Frontiers in evolutionary neuroscience 4, 12, doi:10.3389/fnevo.2012.00012 (2012).

3 Doupe, A. J. & Kuhl, P. K. Birdsong and human speech: common themes and mechanisms. Annu Rev Neurosci 22, 567–631 (1999).

4 Konopka, G. & Roberts, T. F. Animal Models of Speech and Vocal Communication Deficits Associated With Psychiatric Disorders. Biological psychiatry 79, 53–61, doi:10.1016/j.biopsych.2015.07.001 (2016).

5 Gadagkar, V. et al. Dopamine neurons encode performance error in singing birds. Science 354, 1278–1282, doi:10.1126/science.aah6837 (2016).

6 Fee, M. S. & Goldberg, J. H. A hypothesis for basal ganglia-dependent reinforcement learning in the songbird. Neuroscience 198, 152–170, doi:10.1016/j.neuroscience.2011.09.069 (2011).

7 Immelmann, K. in Bird Vocalisations (ed R.A. Hinde) 61–74 (Cambridge University Press, 1969).

8 Price, P. H. Developmental determinants of structure in zebra finch song. J Comp Physiol Psychol 93, 260–277 (1979).

9 Mandelblat-Cerf, Y., Las, L., Denisenko, N. & Fee, M. S. A role for descending auditory cortical projections in songbird vocal learning. Journal Article 3, doi:10.7554/eLife.02152 (2014).

10 Roberts, T. F. et al. Identification of a motor-to-auditory pathway important for vocal learning. Nat Neurosci, doi:10.1038/nn.4563 (2017).

11 Roberts, T. F., Gobes, S. M., Murugan, M., Olveczky, B. P. & Mooney, R. Motor circuits are required to encode a sensory model for imitative learning. Nat Neurosci 15, 1454–1459, doi:10.1038/nn.3206 (2012).

12 London, S. E. & Clayton, D. F. Functional identification of sensory mechanisms required for developmental song learning. Nat Neurosci 11, 579–586, doi:10.1038/nn.2103 (2008).

13 Bottjer, S. W. & Altenau, B. Parallel pathways for vocal learning in basal ganglia of songbirds. Nat Neurosci 13, 153–155, doi:10.1038/nn.2472 (2010).

14 Bottjer, S. W., Miesner, E. A. & Arnold, A. P. Forebrain lesions disrupt development but not maintenance of song in passerine birds. Science 224, 901–903 (1984).

15 Brainard, M. S. & Doupe, A. J. Interruption of a basal ganglia-forebrain circuit prevents plasticity of learned vocalizations. Nature 404, 762–766, doi:10.1038/35008083 (2000).

16 Hoffmann, L. A., Saravanan, V., Wood, A. N., He, L. & Sober, S. J. Dopaminergic Contributions to Vocal Learning. J Neurosci 36, 2176–2189, doi:10.1523/JNEUROSCI.3883-15.2016 (2016).

17 Scharff, C. & Nottebohm, F. A comparative study of the behavioral deficits following lesions of various parts of the zebra finch song system: implications for vocal learning. J Neurosci 11, 2896–2913 (1991).

18 Haesler, S. et al. Incomplete and inaccurate vocal imitation after knockdown of FoxP2 in songbird basal ganglia nucleus Area X. PLoS Biol 5, e321, doi:10.1371/journal.pbio.0050321 (2007).

19 Ali, F. et al. The basal ganglia is necessary for learning spectral, but not temporal, features of birdsong. Neuron 80, 494–506, doi:10.1016/j.neuron.2013.07.049 (2013).

20 Pidoux, L., Leblanc, P. & Leblois, A. A subcortical circuit linking the cerebellum to the basal ganglia engaged in vocal learning. bioRxiv, doi:10.1101/198317 (2017).

21 Keller, G. B. & Hahnloser, R. H. Neural processing of auditory feedback during vocal practice in a songbird. Nature 457, 187–190, doi:10.1038/nature07467 (2009).

22 Schultz, W. Neuronal Reward and Decision Signals: From Theories to Data. Physiological reviews 95, 853–951, doi:10.1152/physrev.00023.2014 (2015).

23 Schultz, W., Dayan, P. & Montague, P. R. A neural substrate of prediction and reward. Science 275, 1593–1599 (1997).

24 Andalman, A. S. & Fee, M. S. A basal ganglia-forebrain circuit in the songbird biases motor output to avoid vocal errors. Proc Natl Acad Sci U S A 106, 12518–12523, doi:10.1073/pnas.0903214106 (2009).

25 Charlesworth, J. D., Warren, T. L. & Brainard, M. S. Covert skill learning in a cortical-basal ganglia circuit. Nature 486, 251–255, doi:10.1038/nature11078 (2012).

26 Tumer, E. C. & Brainard, M. S. Performance variability enables adaptive plasticity of ‘crystallized’ adult birdsong. Nature 450, 1240–1244, doi:10.1038/nature06390 (2007).

27 Yttri, E. A. & Dudman, J. T. Opponent and bidirectional control of movement velocity in the basal ganglia. Nature 533, 402–406, doi:10.1038/nature17639 (2016).

28 Panigrahi, B. et al. Dopamine Is Required for the Neural Representation and Control of Movement Vigor. Cell 162, 1418–1430, doi:10.1016/j.cell.2015.08.014 (2015).

29 Howe, M. W. & Dombeck, D. A. Rapid signalling in distinct dopaminergic axons during locomotion and reward. Nature 535, 505–510, doi:10.1038/nature18942 (2016).

30 Howard, C. D., Li, H., Geddes, C. E. & Jin, X. Dynamic Nigrostriatal Dopamine Biases Action Selection. Neuron 93, 1436–1450 e1438, doi:10.1016/j.neuron.2017.02.029 (2017).

31 Hu, H. Reward and Aversion. Annu Rev Neurosci 39, 297–324, doi:10.1146/annurev-neuro-070815-014106 (2016).

32 Canopoli, A., Herbst, J. A. & Hahnloser, R. H. A higher sensory brain region is involved in reversing reinforcement-induced vocal changes in a songbird. J Neurosci 34, 7018–7026, doi:10.1523/JNEUROSCI.0266-14.2014 (2014).

33 Zann, R. A. The Zebra Finch: a Synthesis of Field and Laboratory Studies. (Oxford University Press, 1996).

34 Williams, H. Birdsong and singing behavior. Ann N Y Acad Sci 1016, 1–30 (2004).

35 Yagishita, S. et al. A critical time window for dopamine actions on the structural plasticity of dendritic spines. Science 345, 1616–1620, doi:10.1126/science.1255514 (2014).

